# LITE-1 Dependent X-Ray Sensitivity in *C. elegans*

**DOI:** 10.1101/766568

**Authors:** Kelli E. Cannon, Meenakshi Ranasinghe, Anhkhoa Do, H. Mariah Pierce, Paul W. Millhouse, Ayona Roychowdhury, Stephen Foulger, Lynn Dobrunz, Jeffrey N. Anker, Mark Bolding

**Affiliations:** Department of Vision Science, University of Alabama at Birmingham School of Optometry, Birmingham, AL; Department of Chemistry, Clemson University, Clemson, SC; Department of Biology, University of Alabama at Birmingham College of Arts and Sciences, Birmingham, AL; Department of Health Professions, University of Alabama at Birmingham School of Medicine, Birmingham, AL; Department of Materials Science, Clemson University, Clemson, SC; Department of Neurobiology, University of Alabama at Birmingham School of Medicine, Birmingham, AL; Department of Radiology, University of Alabama at Birmingham School of Medicine, Birmingham, AL

**Keywords:** *C. elegans*, x-ray, optogenetics, LITE-1

## Abstract

We report the finding that *C. elegans* display X-ray avoidance behavior at high but well tolerated doses, and that this behavior appears to require LITE-1, a gustatory receptor that has been implicated in UV avoidance behavior. We recorded acute behavioral responses of wildtype worms to increasing intensities of X-ray stimulation and found a positive stimulation-response relationship. Mutant strains of worms with dysfunctional photoreceptor proteins LITE-1 and GUR-3 were assayed, and the X-ray avoidance response was found to be nearly absent in LITE-1 mutants but not GUR-3 mutants, suggesting a prominent role for LITE-1 in the detection of X-rays. These findings may be important for developing optogenetics tools to stimulate cells in deep tissue using X-rays, for understanding the mechanism of LITE-1 signaling, and for understanding how organisms may respond to radiation.

## Introduction

Neuroscientists today can manipulate neural activity with unprecedented precision. Spatially restricted genetically targeted cell-type-specific expression of proteins gives researchers the ability to test very specific hypotheses about nervous system function. The best established and most influential technique is optogenetics, which employs light-sensitive photoreceptor proteins to control neuronal activity using light. Since its debut in 2005, optogenetics has lead to many new discoveries, been extensively developed and adapted, and become an indispensable tool in the neuroscientist’s armamentarium. While the transmittance of visible light through brain tissue is high enough to make practicable the use of unaided light to transmit a signal through a few millimeters of tissue (Aravanis et al. 2007), visible photons have low penetrance through the skull, necessitating the still relatively invasive surgical implantation of a fiber optic light source through the skull. In the interest of obviating the craniotomy required by this technique as currently practiced, researchers have begun investigating the use of alternative frequencies of electromagnetic radiation, such as X-rays, which are capable of transmitting through skull and brain tissue with high efficiency.

One method currently under investigation by our group involves radioluminescent nanoparticles (RLPs) delivered to the extracellular space surrounding neurons transgenically expressing photoreceptors.The RLPs absorb X-ray photons and emit visible light photons in a process similar to fluorescence, thereby converting the X-ray signal into a visible light signal that can be detected by transgenic photoreceptors to control neural activity. This technique, referred to as X-ray optogenetics, is expected to require a highly sensitive photoreceptor, as we have found that, at physiologically amenable RLP concentrations and X-ray doses, RLPs emit only a small fraction of the high intensities of light delivered by fiber optic cables in traditional optogenetics setups (unpublished data). Alternatively, a protein that is inherently sensitive to X-rays could potentially bypass the need for RLPs and activate neurons directly.

High-energy photons, including UV, X-rays, and *γ*-rays, can be harmful to life, leading to the generation of reactive oxygen species (ROS) and possible DNA damage. Therefore, quick and effective detection of high energy radiation as noxious and to be avoided can be a highly advantageous trait. The most common form of high-energy photons encountered by organisms is UV light. Extra-ocular light-sensitive proteins such as LITE-1, GR28B and TRP1A have been found to detect high energy UV light in nociceptor neurons of invertebrates (Guntur et al. 2015; Xiang et al. 2010; Birkholz and Beane 2017; Edwards et al. 2008). LITE-1 and GUR-3, both recently discovered in *C. elegans*, function as UV photoreceptors in the worms (Edwards et al. 2008; Bhatla et al. 2015).

Based on reports of the high sensitivity of LITE-1 to UV (Gong et al. 2016), we investigated whether LITE-1 is also sensitive to X-rays by testing the effects of X-ray stimulation on worm behavior. We found that application of X-rays induces a clear increase in activity that is dependent on the intensity of the X-ray stimulation. Importantly, the effect was absent in worms deficient for LITE-1, demonstrating that the X-ray avoidance response requires the activation of LITE-1. Together, these results suggest that LITE-1 might be useful for X-ray optogenetics.

## Methods

### Experimental Model

*C. elegans* nematodes were maintained at 20 °C on NGM agar plates seeded with OP50 *E. coli* lawns (Brenner 1974). All strains of *C. elegans* were obtained from the Caehabditiris Genetics Center, which is funded by the NIH Office of Research Infrastructure Programs (P40 OD010440). We investigated one wildtype strain (N2) and three mutant strains (RB1755, KG1180 and MT21793) to determine if either of two endogenous photoreceptors, LITE-1 or GUR-3, plays a role in the X-ray avoidance response. RB1755 and KG1180 express mutated, dysfunctional GUR-3 and LITE-1, respectively. MT21793 is a double mutant with dysfunctional GUR-3 and LITE-1 that has severely defective responses to light (Bhatla and Horvitz 2015).

### X-ray stimulation

X-rays were generated by a iMOXS-MFR tungsten target X-ray unit with polycapillary optics (XOS) to focus the X-ray beam to a FWHM diameter of 0.85 mm at the level of the agar surface, located approximately 5 cm from the tip of the capillary attachment. The unit was operated at 50kV and the current was varied from 0 to 600 *µ*A to achieve different stimulation intensities. At the highest intensity, a radiation dose of 1 Gy/s was approximated using RADSticker dosimeter stickers (JP Laboratories, Inc.). The dose decreased to approximately 0.2 Gy/s when the X-ray unit was operated at 25% of the maximum current, i.e., 150 *µ*A. Although the exact X-ray doses may vary somewhat from these values, we expect these dose estimates to be accurate within a factor of two. An internal X-ray shutter was used to control the timing of the X-ray stimulation.

### Imaging Setup and Calibration

Worm behavioral responses to X-ray stimulation were recorded on a custom imaging setup, as shown in Figure 1a,b. Video data was recorded using an Amscope MU1003 10MP CMOS camera with a 0.7X-180X magnification lens. Back lighting was provided by a white LED source with diffuser. The X-ray unit was mounted above the stage.The stage was positioned using a motorized x-y translator controlled by the joystick of an Xbox controller (Microsoft, Redmond WA). The entire imaging setup was enclosed in a lead box.

**Fig. 1.**
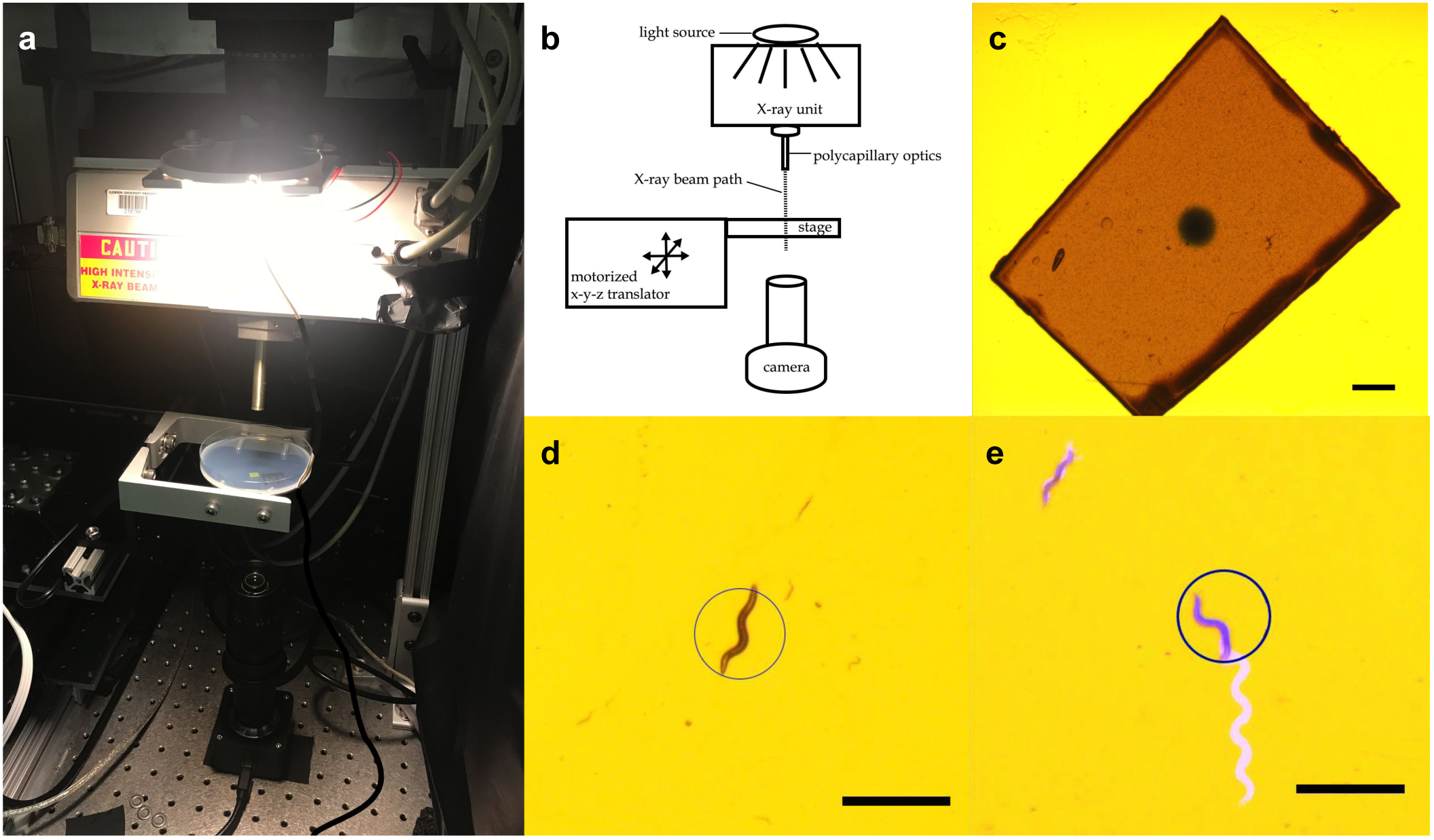
Experimental Setup. a) photograph of setup b) diagram of setup c) X-ray spot visualized with a piece of radiochromic film placed on the agar surface. d) Irradiation spot is represented as a circle overlayed on a worm to show stimulation targeting. e) An example of a positive X-ray avoidance response, where the worm’s movement over the duration of the 10 second X-ray pulse is shown in white. Scale bars indicate 1 mm. duration of the 10 second X-ray pulse is shown in white. Scale bars indicate 1 mm.

The irradiation zone—a sharply defined, 0.85 mm diameter spot on the agar surface—was visualized using a small piece of radiochromic film (Gafchromic, XR-QA2) placed on the agar surface (see Figure 1c). The camera recording software (Amscope 3.7) was used to annotate the video monitor with a circle outlining the irradiation zone as visualized with the gafchromic film for targeting purposes.

X-ray shutter timing was synced to video timing post hoc using a second video recording device outside of the lead box that acquired audio data. The camcorder collected video of the live recording feed with a timestamp displayed as well as audio data capturing the click of the X-ray shutter opening and closing. The timestamp on the video recording updated at a rate of 1 Hz, limiting the precision of the timing data to ±1 s.

### Experimental Design

All experiments were conducted on OP50-lawned 100 mm NGM plates inoculated with approximately 20-50 worms 24-48 hours prior. Looking at one worm per trial, a total of 130 trials were conducted on worms from 11 different plates, targeting 3 to 20 individual worms from each plate. A small (*∼*1 cm^2^) piece of gafchromic film was placed on the agar surface of each plate before the plate was placed, uncovered, on the imaging stage in the irradiation chamber. After calibrating the location of the X-ray spot with the gafchromic film as described above, an adult hermaphrodite was selected and the stage was moved to position the worm in the center of calibrated X-ray spot. Each trial began with 30 seconds during which baseline behavior was recorded with the X-ray shutter closed. After 30 seconds, if the worm moved out of the center of the calibrated spot, the stage was moved to reposition the worm in the irradiation zone. Once the worm was in position, the shutter was opened to deliver a 10 second X-ray pulse. The pulse was terminated by closing the shutter, and worm behavior was recorded for another 10 seconds after stimulation offset.

### Stimulation-response Experiments

Behavioral responses of wildtype (N2) worms to increasing intensities of X-ray stimulation were recorded. Five X-ray intensities between zero and the maximum intensity deliverable by our iMOX X-ray unit, were tested. Intensities of approximately 0, 0.2, 0.5, 0.7, and 1.0 Gy/s were achieved by setting the current applied to the X-ray tube to 0, 150, 300, 450, and 600 *µ*A, respectively. Twenty trials of each the null and highest intensities, and ten trials at each of the three intermediate intensities were collected, for a total of 70 trials. The five intensities were delivered in an interspersed and randomized order.

### Strain Comparison Experiments

The responses of three mutant strains of *C. elegans*, RB1755, KG1180 and MT21793, were tested at the null and maximum X-ray intensities, as described above. Ten trials at each of the two intensities were conducted for each strain for a total of 60 trials.

### Data Analysis

Video recordings were stabilized in Matlab (due to plate movement from recentering the target worm in the irradiation zone) and analyzed using Worm Lab software (version 2019.1.1) to track and measure the activity levels of worm subjects. Prior analysis of an independent preliminary dataset (unpublished data) found the behavioral metric most sensitive to the X-ray avoidance response was activity, in units of body area per minute, as defined by CeleST tracking software (Restif et al. 2014). Activity time courses were combined with stimulation timing info in Matlab (R2019a) to calculate the peak change in activity—the maximum activity reached during the 10-second X-ray pulse minus the average activity during 30-second baseline period—normalized to the baseline activity for each trial. We defined a positive avoidance response as an increase in activity greater than 45%, using the largest increase observed in the negative control condition to guide threshold selection. Three trials were excluded due to tracking problems. Figure 1e illustrates a typical positive avoidance response, wherein the worm increases forward locomotion to escape the irradiation zone, which is indicated by a blue circle. This image was created by calculating the difference between the maximum and minimum intensities of each pixel across all frames during the 10 second X-ray pulse and adding the difference image to the blue component of the RGB image of the first frame.

## Results

### wildtype worms respond to X-rays in a intensity-dependent manner

To probe the sensitivity of wildtype worms to increasing intensities of X-ray stimulation, we operated the X-ray unit at a range of currents and recorded behavioral responses to the different X-ray intensities. Worm activity showed a marked increase during X-ray stimulation when the X-ray was operated at the unit’s maximum current, 600 mA (Figure 2b,c,d). The increase in activity during X-ray stimulation was less prevalent at the 0.5 and 0.7 Gy/s intensity conditions (Figure 2c,d). Worm activity appeared unaltered by X-ray stimulation at either 0.2 Gy/s or the null intensity condition (Figure 2a,c,d). The average peak change in activity displayed a positive correlation to X-ray intensity, as shown in Figure 2c. The fraction of worms displaying an avoidance response—i.e., a >45% increase in activity—was zero for both the null and 0.2 Gy/s intensity conditions (Figure 2d). At 0.5 and 0.7 Gy/s X-ray intensities, half the worms exhibited a response. At the highest intensity tested, 70% of wildtype worms displayed an avoidance response.

**Fig. 2.**
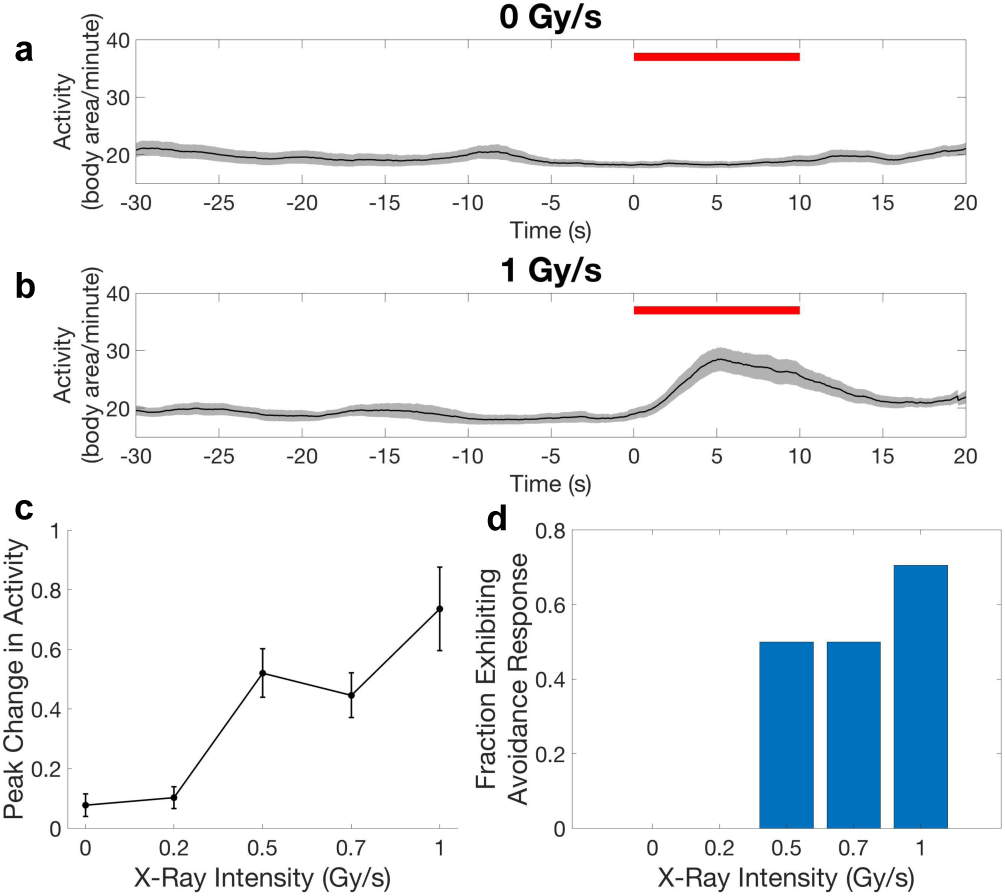
Stimulation-response relationship of wildtype worms. a,b) Activity time courses averaged across worms at 0 and 1 Gy/s stimulation intensity conditions. Shading indicates standard error of the mean time series. X-ray stimulation timing is indicated by red bar. c) The mean peak change in activity normalized to baseline activity increases in response to increasing stimulation intensities. Error bars are standard error. d) The fraction of worms exhibiting an X-ray avoidance response increases at increasing stimulation intensities.

### Worms with dysfunctional photoreceptor proteins have a dysfunctional X-ray avoidance response

We next tested whether either of two *C. elegans* photoreceptor proteins, LITE-1 or GUR-3, plays a role in the behavioral response to X-rays. We subjected three mutant strains of *C. elegans* that exhibit dysfunctional photoresponses due to mutations in these proteins (Bhatla and Horvitz 2015) to the null and the maximum X-ray intensity conditions. Activity levels of RB1755, the GUR-3 only mutant, increased in response to X-ray stimulation (Figure 3b), similarly to the wild type strain. In contrast, the activities of KG1180, the LITE-1 only mutant, and MT21973, the double mutant, appeared largely unaltered by stimulation (Figure 3c,d). The average peak change in activity showed a significant increase from the null to the maximum intensity conditions for the N2 and RB1755 strains (N2 t(16) = -4.68, p = 0.00025; RB1755 t(9) = -5.37, p = 0.00045; Figure 3e). No significant difference between the two conditions was observed for the KG1180 or MT21973 strains (KG1180 t(9) = -0.59, p = 0.57; MT21973 t(9) = - 1.10, p = 0.30; Figure 3e). The fraction of worms exhibiting an avoidance response increased from zero to 0.9 between the null and maximum intensity conditions for the GUR-3 only mutant strain (Figure 3f). No significant increase in the fraction of worms displaying avoidance behavior was observed for either of the LITE-1 mutant strains.

**Fig. 3.**
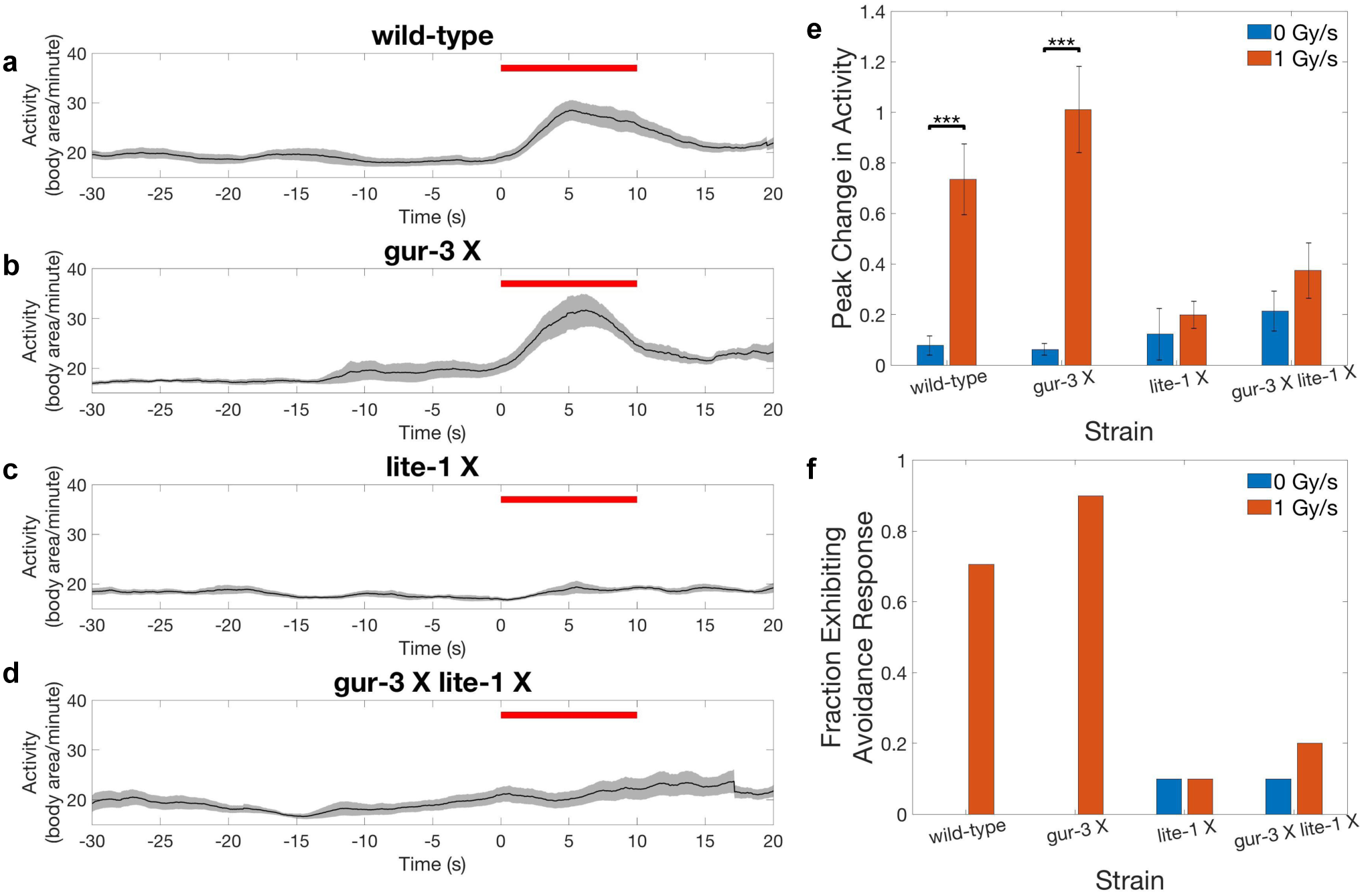
Mutant strains lacking functional LITE-1 have a defective X-ray avoidance response. a-d) Activity time courses averaged across worms of the wildtype strain and each of the three mutant strains for the 1 Gy/s X-ray stimulation intensity condition. Shading indicates standard error of the mean time series. X-ray stimulation timing is indicated by red bar. e) The mean peak change in activity normalized to baseline activity in response to 0 Gy/s and 1 Gy/s X-ray stimulation intensities for each strain. Error bars are standard error. *** indicates a significance level <0.001. f) The fraction of worms exhibiting a positive avoidance response to 0 Gy/s and 1 Gy/s X-ray intensities for each of the four strains.

## Discussion

Here we show that wildtype *C. elegans* exhibit a locomotory avoidance response to X-ray stimulation that is dependent on LITE-1.

The X-ray avoidance response was observed in some wildtype worms at the median X-ray intensity of 0.5 Gy/s, but it was not seen at the lower intensity of 0.2 Gy/s, suggesting the threshold X-ray intensity needed to trigger the response may lie between 0.2 - 0.5 Gy/s. While the response appeared intact in the GUR-3 null strain, it was severely deficient in the two strains lacking functional LITE-1. LITE-1 and GUR-3 share 40% homology and are both involved in the inhibition of feeding response to UV light (Bhatla and Horvitz 2015; Gong et al. 2016); however, only LITE-1 is thought to participate in the avoidance response, as the response appears intact in GUR-3 mutants (Bhatla and Horvitz 2015). Therefore, our finding that X-ray avoidance also appears intact in the strain with mutated GUR-3, but functional LITE-1, is in line with previous findings demonstrating a prominent role for LITE-1, but not GUR-3, in locomotory avoidance behavior. The possibility that GUR-3 is similarly sensitive to X-rays remains open.

This study has two important limitations: the consistency of X-ray targeting and stimulation and the temporal precision with which stimulation could be synced to behavior. Targeting of the X-ray stimulation varied from trial to trial due to 1) slight variation in targeting at the start of the 10 s pulse due to the limited stage positioning accuracy afforded by Xbox controller joystick, and 2) variation in worm movement during positioning and stimulation, leading to variability in the duration of stimulation to which each worm was exposed, with fast moving worms spending less time in the irradiation zone compared to slower worms. Therefore, the stimulation-response experiments performed here using different intensities of X-ray stimulation only approximate the underlying dose-response relationship, as X-ray dose is a function of intensity over time. Variability in worm movement also affected the timing of the stimulation, as sometimes the spot was positioned in the predicted path of more quickly moving worms, such that the worm passed through the spot up to a few seconds after the shutter opened. This, along with the relatively low precision with which the timing of X-ray stimulation could be synced to behavior (on the order of ±1 s), limits our analysis of the temporal aspects of the response. Further studies are needed to better elucidate the dose-response relationship and timing of the *C. elegans* X-ray avoidance response. Despite these limitations, our experiments clearly demonstrate that X-rays induce an avoidance behavior in worms that is dependent upon X-ray intensity and requires LITE-1.

X-ray detection by ocular photoreceptor proteins such as rhodopsin has been suggested by reports of X-ray phosphenes (i.e., sensations of light produced by X-rays) experienced by astronauts and radiation-therapy patients (Fuglesang et al. 2006; Lipetz 1955; Thariat et al. 2016). In addition, there have been numerous reports of behavioral and electroretinogram (ERG) responses to X-rays in animal models (Bachofer and Wittry 1961; Baldwin and Sutherland 1965; Baily and Noell 1958; Martinsen and Kimeldorf 1972). However, LITE-1, which more closely resembles a family of gustatory chemoreceptors than any known photoreceptor, appears to operate by a different mechanism in the detection of light. Critically, it lacks the prototypical light-absorbing small-molecule chromophore partner (e.g., retinal) seen in other photoreceptor complexes such as opsins and cryptochromes. Consistent with its chemoreceptor lineage, it has been suggested that LITE-1 detects not UV photons themselves, but rather a reactive oxygen species (ROS) photoproduct, such as H_2_O_2_, which is generated by UV light interacting with photosensitizing molecules such as riboflavin (Bhatla and Horvitz 2015). In support of hypothesis, wildtype worms have been found to exhibit an avoidance response to H_2_O_2_, and this response is nearly absent in LITE-1 mutants (Bhatla and Horvitz 2015). X-rays can also generate oxygen radicals by interacting with any available source of oxygen, including water (Le Caër 2011). Thus, it is plausible that an ROS intermediary may be involved in the mechanism underlying LITE-1’s sensitivity to X-rays.

As *C. elegans* display robust thermotaxic behavior and are capable of detecting changes in temperature as small as 0.05 °C (Clark et al. 2006), control experiments were important to ensure that the *C. elegans* UV avoidance response was not the result of temperature changes induced by the light. The maximum increases in temperature from 10 s of exposure at maximum 1 Gy dose rate (i.e., 10 J/kg cumulative dose after 10 s), conservatively based only on the heat capacity of water (4.2 kJ/kg/K), is 2.4 mK, 20-fold less than 0.05 °C. In our experiment, the worms responded within 2 s of exposurem (just 2 Gy cumulative dose) and the heat loss from thermal diffusion and convection would have further decreased the temperature increase in the irradiated region. As a result, we expect any thermal effects to be negligible. Additionally, the observed absence of an avoidance response in the two LITE-1 mutant strains demonstrates that any heat generated by the X-rays in these experiments has a negligible effect on worm activity without functional LITE-1. As LITE-1 itself is not known to have thermoreceptor properties, it is unlikely that temperature changes played a role in the results.

If LITE-1 is activated by X-rays (or a chemical product generated by X-rays) alone, intermediary RLPs to transduce the X-ray signal into a UV light signal may be unnecessary for X-ray optogenetics, depending on the efficiency with which X-rays activate LITE-1. It is plausible that adding the RLPs could enhance LITE-1 activation by X-rays, either by converting some of the X-rays to UV light, which may more effectively activate LITE-1, or by scattering the X-rays, thereby increasing the probability of interaction with LITE-1. Important to keep in mind when employing ionizing radiation, the radiation dose is best kept to a minimum to avoid acute radiation sickness and off-target effects. Oxidative stress resulting from unchecked ROS production is undoubtedly deleterious to cells at very high levels and/or long exposures. However, cells naturally generate ROS as a byproduct of aerobic energy metabolism, and a regiment of antioxidant mechanisms have evolved to counterbalance their production and mitigate harmful effects. There is even some evidence that ROS such as H_2_O_2_ can have a hormetic effect, such that low doses can have opposite, beneficial effects, increasing resistance to oxidative stress, life span, and sensory function (Schulz et al. 2007; Ristow and Schmeisser 2011; Li et al. 2016). It is plausible that short pulses of low intensity X-rays could generate acute increases in ROS levels capable of being detected by a highly sensitive protein, but insufficient to overwhelm the cell’s natural antioxidant systems. Key to harnessing the advantages of this technology will be maximizing the efficiency of signal transmission in order to minimize off-target effects of radiation. Further investigation is needed to assess the efficiency with which LITE-1 can be activated by X-rays compared UV and to determine whether adding RLPs will decrease the threshold radiation dose needed to activate LITE-1.

In sum, we found that the *C. elegans* UV photoreceptor LITE-1 is sensitive to X-ray stimulation, as demonstrated by a conspicuous X-ray avoidance response seen only in worms with intact LITE-1. This suggests that LITE-1 may be a useful photoreceptor for X-ray optogenetics, enabling low doses of X-rays to replace invasive fiber optic implants for highly precise manipulation of neural activity.

## ACKNOWLEDGEMENTS

Funding for this study was provided by NSF EPSCoR OIA-1632881.

